# Switch-like control of helicase processivity by single-stranded DNA binding protein

**DOI:** 10.1101/2020.07.09.194779

**Authors:** Barbara Stekas, Masayoshi Honda, Maria Spies, Yann R. Chemla

**Affiliations:** Department of Physics, University of Illinois, Urbana-Champaign, Urbana; Department of Biochemistry, Carver College of Medicine, University of Iowa, Iowa City; Center for the Physics of Living Cells, University of Illinois, Urbana-Champaign, Urbana

## Abstract

Helicases utilize the energy of NTP hydrolysis to translocate along single-stranded nucleic acids (NA) and unwind the duplex. In the cell, helicases function in the context of other NA-associated proteins which regulate helicase function. For example, single-stranded DNA binding proteins are known to enhance helicase activity, although the underlying mechanisms remain largely unknown. *F. acidarmanus* XPD helicase serves as a model for understanding the molecular mechanisms of Superfamily 2B helicases, and previous work has shown that its activity is enhanced by the cognate single-stranded DNA binding protein RPA2. Here, single-molecule optical trap measurements of the unwinding activity of a single XPD helicase in the presence of RPA2 reveal a mechanism in which XPD interconverts between two states with different processivities and transient RPA2 interactions stabilize the more processive state, activating a latent “processivity switch” in XPD. These findings provide new insights on mechanisms of helicase regulation by accessory proteins.

## Introduction

Helicases are molecular machines that use the energy of NTP hydrolysis to separate the strands of nucleic acid (NA) duplexes. A large class of helicases unwind by translocating directionally along a single NA strand and can utilize this translocation activity to displace NA-bound proteins (1–4). In the cell, helicases are involved in many essential genome maintenance processes, including replication, recombination, and repair (5–7). In a number of instances, the same helicase is known to carry out several of these disparate functions (4,8,9). Thus, helicase activity must be tightly regulated, not only to prevent indiscriminate unwinding of duplex DNA in the cell, which could lead to genome instability, but also to define the context-dependent role of the helicase. How this regulation occurs is often unclear, but growing evidence points to interactions with protein partners as one regulatory mechanism (4,8,9).

Xeroderma pigmentosum group D (XPD) protein is a 5’ to 3’ DNA helicase that serves as a model for understanding members of the structural Superfamily 2B (SF2B) of helicases, a group that includes yeast Rad3 and human FANCJ, RTEL, and CHLR1 (9–12). Human XPD is part of transcription factor IIH (TFIIH) and plays a vital role in nucleotide excision repair (NER) (13–16). It has also been shown to participate in chromosome segregation (17) and in the cell’s defense against retroviral infection (18). Structurally, XPD consists of a conserved motor core (**Fig. 1a**)—helicase domains 1 and 2 (HD1 and HD2)—that couples ATP binding and hydrolysis to translocation on ssDNA, a FeS cluster-containing domain, and a unique ARCH domain that encircles the translocating strand (19–23).

**Figure 1.**
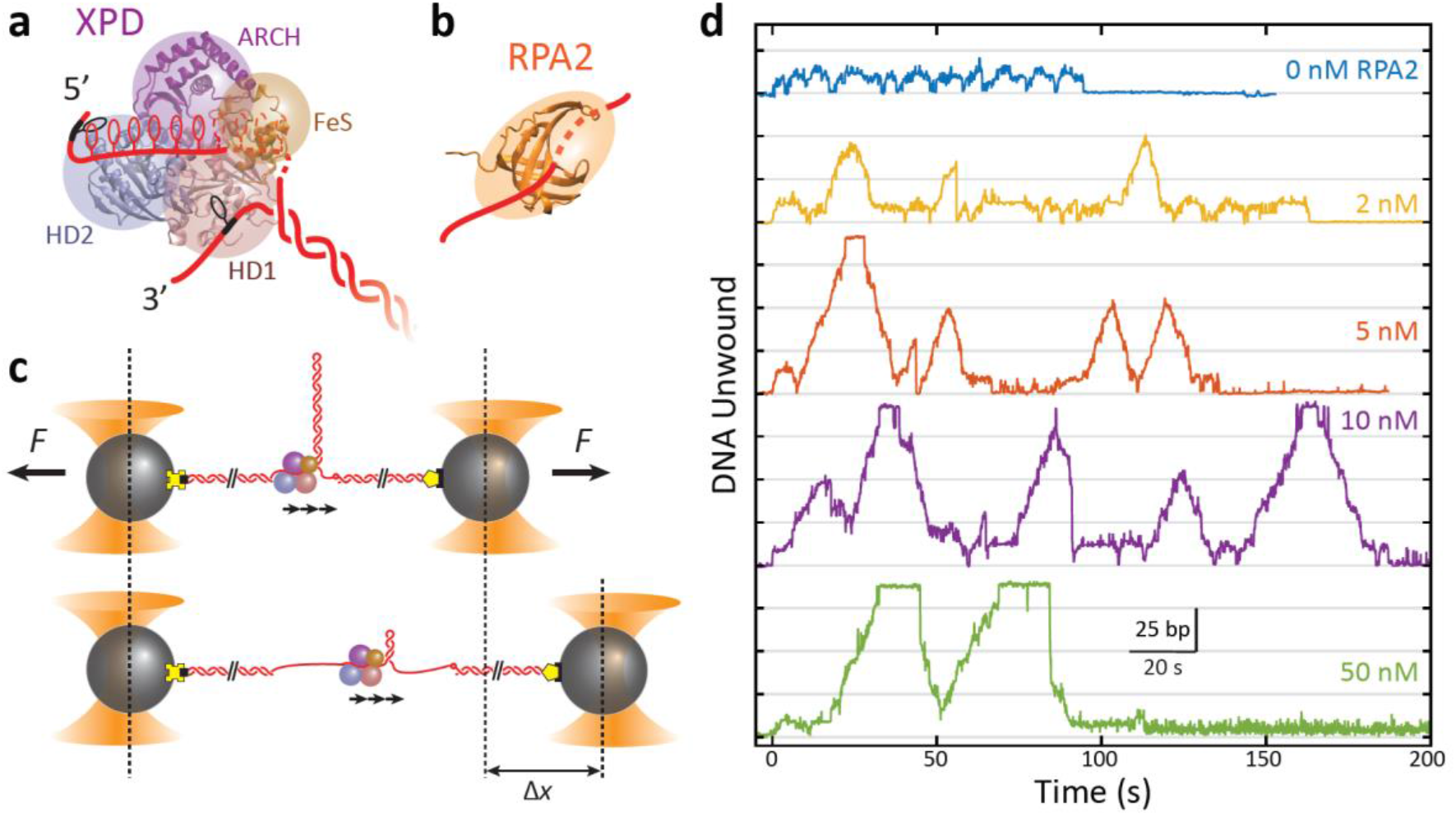
RPA2 increases XPD helicase processivity. **a** Schematic of *Fac*XPD based on *S. acidocaldarius* XPD structure (PDB 3CRV). XPD is composed of four domains: Helicase Domain 1 and 2 (HD1, blue; HD2, pink) which form the motor core, and the ARCH domain (purple) and FeS cluster (brown). A model of the DNA fork (red) shows ~10 nt bound to the motor core and secondary contacts (black) to each strand of the fork. **b** Schematic of *Fac*RPA2 based on partial crystal structure of *M. maripaludis* RPA (PDB 2K5V). RPA2 consists of a single OB fold that binds to ssDNA in the C-shaped cavity. **c** Single-molecule hairpin unwinding assay. A DNA hairpin consisting of an 89-bp stem and (dT)_4_ loop is tethered between trapped beads by biotin-streptavidin (yellow cross) and digoxigenin-antibody linkages (pentagon) and held at a constant force. A 10-nt poly-dT site at the 5’ end of the hairpin allows one XPD molecule to load, and an abasic site at the 3’ end prevents XPD unwinding past the hairpin. Unwinding of the hairpin by XPD increases the end-to-end extension of the construct by an amount, *Δx*, proportional to the number of base pairs unwound. Arrows indicate the 5’ to 3’ direction of XPD translocation along ssDNA. **d** Representative traces of a single molecule of XPD unwinding in the presence of varying concentrations of RPA2 (0-50 nM) at constant force (*F* = 12 pN). ATP and RPA2 are added at *t* = 0 s. XPD processivity increases with RPA2 concentration.

As it functions on ssDNA, XPD is expected to encounter other proteins such as Replication protein A (RPA), which binds single-stranded DNA non-sequence specifically. Single-stranded DNA binding proteins function as protectors of ssDNA against nucleolytic damage and play important regulatory roles (24). Due to their ubiquity, frequent interactions with other DNA-bound proteins are likely (25). Single-stranded binding proteins are known to stimulate the DNA unwinding activity of SF2 helicases, although the mechanisms of enhancement remain elusive (26–31). In the present work, we investigate the effect of single-stranded DNA binding proteins on helicase unwinding activity using XPD from *Ferroplasma acidarmanus* as a model system due to the availability of biochemical (31) and single-molecule kinetic data (32–34), as well as structural information from homologs (20–22,35). Prior work (31) has shown that *Fac*XPD unwinding is enhanced greatly by one type of cognate ssDNA-binding protein in *F. acidarmanus*, RPA2, more so than a second cognate protein, RPA1, or by heterologous proteins. *Fac*RPA2 has a simple, single-domain architecture with one OB fold that occludes ~4 nt of ssDNA (**Fig. 1b**) (31). Single-molecule experiments demonstrated that XPD can occupy the same DNA strand as RPA2 and bypass RPA2-bound ssDNA (32). Crucially, no direct, specific interactions between RPA2 and XPD in solution have been observed (31).

How an ssDNA-binding protein with no known protein-protein contacts to the helicase stimulates helicase-mediated DNA unwinding activity remains an open question, and makes *Fac*XPD and *Fac*RPA2 an intriguing system for studying the stimulatory effects of ssDNA-binding proteins on helicases. Mechanisms for enhancement of unwinding activity can loosely be placed in three categories (31). First, an ssDNA-binding protein could destabilize the DNA duplex at the ssDNA-dsDNA junction ahead of the helicase, facilitating its motion forward. Second, it could enhance unwinding by sequestering ssDNA and providing a physical barrier to helicase backsliding, i.e. rectifying helicase forward motion. Lastly, it could activate the helicase for processive unwinding through direct interactions with the helicase-DNA complex.

Here, we use a single-molecule optical trap assay to observe individual proteins of XPD unwinding DNA and to analyze the effects of RPA2 on their unwinding activity. While previous reports have shown that RPA2 is able to enhance XPD activity (31,32), they have not defined what aspects of activity is enhanced or provided a mechanism. Our results show that RPA2 primarily increases XPD processivity, or the maximum number of base pairs unwound. XPD exhibits repeated attempts to unwind duplex DNA and RPA2 increases the frequency of attempts that have high processivity. While RPA2 can transiently destabilize duplex DNA, this does not promote XPD unwinding. Instead, data point to a mechanism in which XPD possesses a latent processivity “switch” which is activated by RPA2. Our measurements shed new light on the mechanisms by which accessory proteins such as single-stranded DNA binding proteins can enhance helicase activity.

## Results

### XPD processivity increases in the presence of RPA2

As shown in **Fig. 1c**, we monitored the activity of a single XPD in the absence and presence of RPA2 in solution from the unwinding of an 89-bp DNA hairpin (**Figure 1—Figure Supplement 1**) tethered between two optically trapped beads and stretched under constant force (12 pN), as described previously (33). Data were collected at a rate of 89 Hz. Unwinding of the hairpin was detected from the increase in the end-to-end extension of the DNA tether, as each broken base pair released 2 nucleotides (see **Methods**). An abasic site positioned at the 3’ end of the hairpin prevented XPD from unwinding the long (1.5-kb) dsDNA handle used to separate the hairpin from the trapped beads (33) (**Figure 1—Figure Supplement 1**). To control the loading of XPD and RPA2 onto the hairpin, we used a custom flow chamber consisting of parallel laminar-flow streams containing different buffers, as described previously (36) (see **Methods**; **Figure 1—Figure Supplement 2**). Placing the hairpin in one stream containing XPD but no ATP for ~1 min., the protein was allowed to bind to a 10-dT ssDNA loading site—approximately equal to the footprint of a single XPD (33,37)—at the 5’ end of the hairpin (**Figure 1—Figure Supplement 1**). Unwinding was initiated by moving the hairpin to a stream containing saturating ATP (500 μM) and varying concentrations of RPA2 (0-50 nM). This procedure ensured that a single XPD was loaded at one time, and that the unwinding observed resulted from that of a single XPD helicase in the absence or presence of many RPA2 molecules in solution (see **Methods**).

**Figure 1d** shows representative traces of a single XPD protein unwinding in increasing [RPA2]. As previously reported, XPD unwound in repeated “bursts,” comprising cycles of forward unwinding motion followed by backward rezipping (33), repeating a number of times (5 ± 1) until the protein dissociated. In the absence of RPA2, we observe that the processivity—the maximum hairpin position reached by XPD during any one burst—was low, on average 15-20 out of 89 bp, consistent with our prior work (33). With increasing [RPA2], XPD unwound farther into the hairpin, occasionally unwinding the entire 89-bp hairpin stem (e.g. *t* = 20 s for 5 nM RPA2 or *t* = 30 s, 70 s for 50 nM RPA2 in **Fig. 1d**).

Each burst represents an attempt by one XPD molecule to unwind the hairpin, and we found that processivity varied greatly from burst to burst. Each burst consisted of varying extents of duplex unwinding followed by rezipping, often to the base of the hairpin. As shown in the representative trace in **Fig. 2a**, several molecular events comprised backward motion: XPD backstepping or backsliding via temporary disengagement from the unwinding DNA strand (**Fig. 2b**: diagram 1a and 1b), or translocation on the other hairpin strand away from the fork junction (2 and 3), all leading to DNA rezipping under its regression force. An example of the latter occurred when XPD completely unwound the hairpin stem and translocated past the dT tetraloop cap onto the opposing strand, allowing the stem to rezip gradually in the protein’s wake (**Fig. 2b**: diagram 2). In a contrasting example, we observed gradual rezipping of the duplex mid-hairpin, which we attributed to XPD disengaging from its DNA strand to switch to the opposing strand (**Fig. 2b**: diagram 3), a behavior reported for other helicases (38,39). This interpretation is corroborated by the observation that gradual rezipping was typically followed by a stall at ~10 bp (**Fig. 2a**, *t* = 58 s) the size of XPD’s footprint, consistent with the protein stalling at the abasic site on the 3’ strand at the base of the hairpin. Subsequent unwinding bursts were presumably due to XPD strand-switching back to the original strand. In contrast, rapid rezipping events due to backsliding usually occurred to the base of the hairpin (**Fig. 2a**, *t* = 44 s), consistent with the protein remaining on the hairpin 5’ strand.

**Figure 2.**
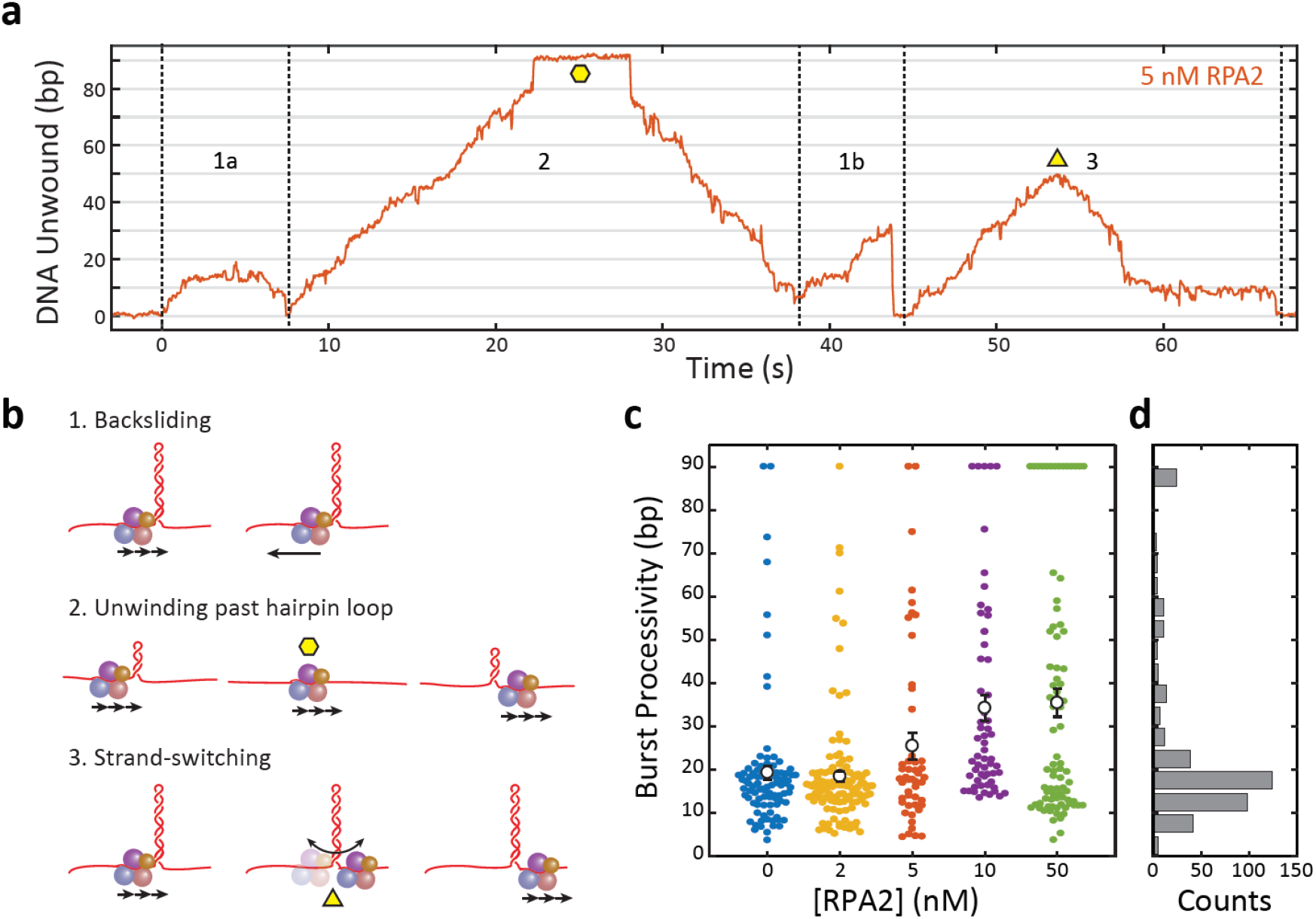
XPD unwinds in bursts of varying processivity whose average increases with RPA2 concentration. **a** Representative trace of a single molecule of XPD unwinding in the presence of RPA2 (5 nM). One XPD exhibits repetitive bursts of activity, making multiple attempts to unwind hairpin DNA. Processivity can vary widely from burst to burst. Example time trace with: (1a) 20 bp- and (1b) 30 bp-processivity bursts composed of forward unwinding followed by backsliding to the hairpin base; (2) a high-processivity burst during which XPD completely unwinds the 90-bp hairpin past the hairpin loop (time point indicated by yellow hexagon) and translocates on the opposing strand, allowing the hairpin to rezip; and (3) a 50 bp-processivity burst during which XPD unwinds and switches strand mid-hairpin (indicated by yellow triangle), allowing the hairpin to rezip. **b** Schematics representing the behaviors in (a). **c** Processivity of each burst (colored circles) vs. RPA2 concentration. The mean processivity (open circles) increases with RPA2 concentration. Error bars represent s.e.m. **d** Histogram of all burst processivities at all RPA2 concentrations.

We analyzed the processivity of each burst as a function of [RPA2], as shown in the scatter plot in **Fig. 2c**. In the absence of RPA2, nearly all the individual burst processivities cluster below 25 bp, with a small fraction (<10%) extending further. When unwinding in the presence of RPA2, the average burst processivity increases with [RPA2] up to 35 bp at 50 nM RPA2. However, the distributions of burst processivity at each RPA2 concentration show a significant fraction at relatively low processivity, ≲25 bp, persisting across all concentrations. As [RPA2] increases, an increasing fraction of bursts fall into a higher processivity tail of the distribution, which includes many cases of complete unwinding of the 89-bp hairpin. The distribution of burst processivity for all [RPA2] (**Fig. 2d**) shows a large population of bursts with average processivity of 15-20 bp, followed by a long tail with processivity extending from 25 to 89 bp.

### Melting of the hairpin by RPA2 does not aid XPD unwinding

We next asked by what mechanism RPA2 increases the processivity of XPD. One possibility is that RPA2 helps destabilize the duplex, assisting XPD in unwinding. In such a duplex melting mechanism (31), RPA2 would presumably bind at the ssDNA-dsDNA junction and melt several base pairs ahead of XPD. Because XPD relies heavily on thermal fluctuations to help it break the base pairing bonds ahead of it (33), RPA2 lowering the energy barrier to unzipping the duplex could plausibly enhance XPD’s unwinding activity.

As shown in **Fig. 3a**, RPA2 is capable of transiently destabilizing hairpin DNA under force of 12 pN. In experiments with RPA2 but without XPD, short melting events are observed, corresponding to the hairpin opening by ~5-6 bp, comparable to the footprint of RPA2 (31), and reannealing shortly thereafter (~20 ms). [Such transient melting is also observed in **Fig. 1d** after XPD dissociation (e.g. at *t* > 110 s for 50 nM RPA2).] These melting events become more frequent as the concentration of RPA2 is increased (**Figure 3—Figure Supplement 1**), in support of melting being RPA2-mediated. Analyzing the times between melting events, we estimate an effective second-order rate constant of (1.2 ± 0.3) × 10^8^ M^-1^ s^-1^ for binding.

**Figure 3.**
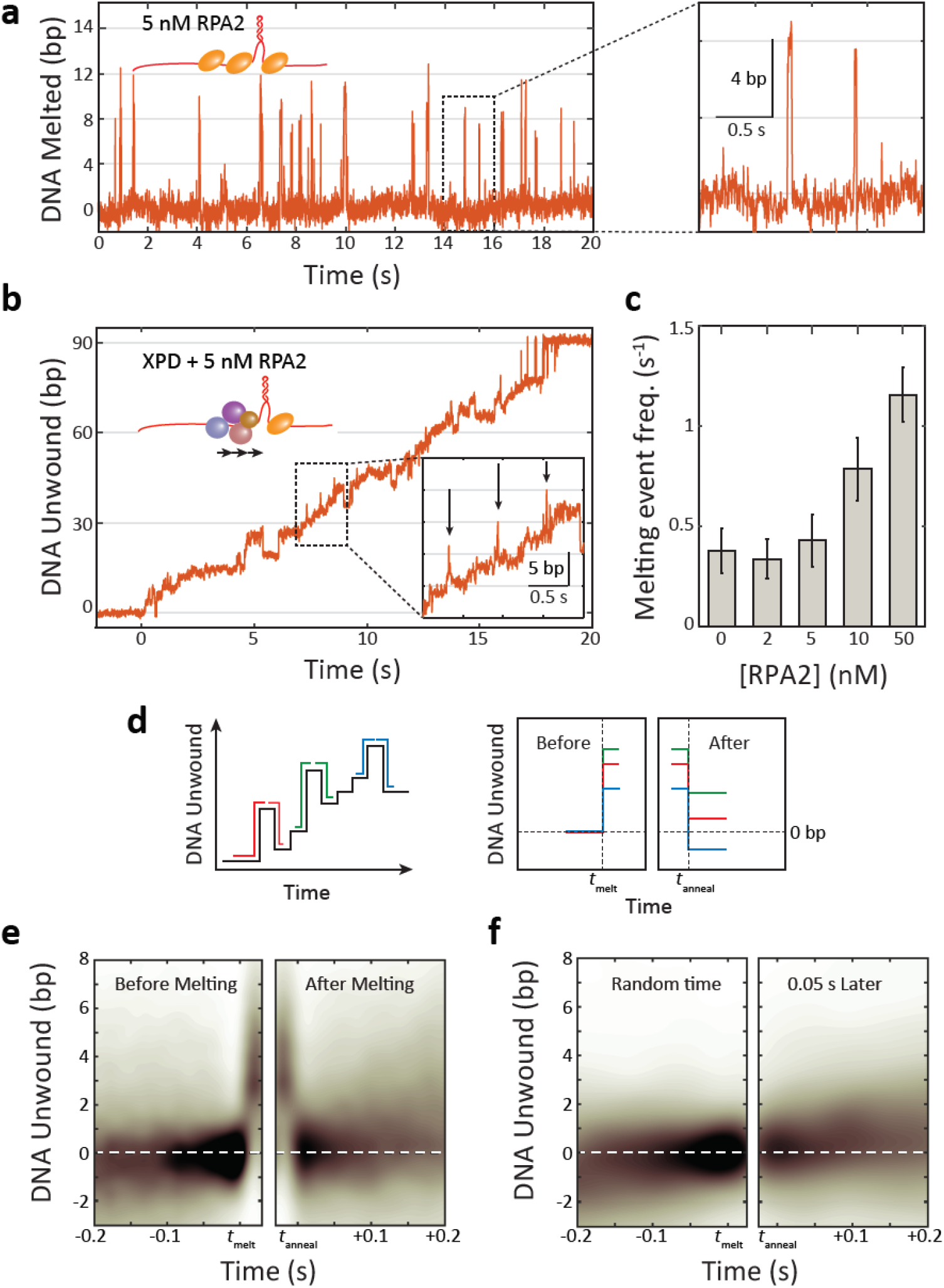
RPA2 transiently melts hairpin duplex but does not assist XPD unwinding. **a** Representative time trace of RPA2 transiently destabilizing hairpin dsDNA at a constant force (*F* = 12 pN). Inset: RPA2 melts ~8 bp, which then rapidly reanneals (see Figure 3—Figure Supplement 1). **b** RPA2 is able to melt dsDNA at a fork occupied with XPD. Inset: transient RPA2-like melting events during XPD unwinding (see Figure 3—Figure Supplement 2). **c** The frequency of RPA2 melting events increases with RPA2 concentration. Error bars represent s.e.m. **d** Analysis of RPA2 melting events and their effect on XPD unwinding. The schematic shows all RPA2 melting events identified and divided into “melting” and “reannealing” transitions. Each transition is aligned temporally such that all melting transitions begin at the same time *t_melt_* and corresponding reannealing transitions end at the same time *t_anneal_*. Both types of transitions are aligned spatially relative to the starting extension before melting (*x* = 0). **e** Probability distribution of all aligned RPA2 melting events. Although RPA2 melts an average of 5 bp of hairpin DNA (left), it reanneals by the same amount (right), and there is no net progress of XPD due to melting. **f** Probability distribution at a random time point (left) and 0.06 s later (right). Net progress of XPD after an RPA2 melting event is indistinguishable from net progress at a random time point. Probability distributions were obtained using kernel density estimation (see Methods).

RPA2-mediated melting events are also observable as XPD unwinds in the presence of RPA2 (**Fig. 3b**). Such events are identifiable over XPD unwinding dynamics due to their characteristic size (~5-6 bp) and short lifetime (~20 ms) (**Fig. 3b**, inset; see **Methods** and **Figure 3—Figure Supplement 2**). While some false positives are detected in the absence of RPA2, they are uncommon and the frequency of detected melting events increases with [RPA2] (**Fig. 3c**), confirming their connection to RPA2 activity. This finding indicates that RPA2 is capable of binding at the same fork junction as XPD and melting DNA ahead of the helicase. However the question remains whether RPA2 transient melting aids XPD in unwinding.

To answer this question, we determined how RPA2-mediated melting events impact the forward progress of the helicase. If duplex destabilization aids in unwinding, we suspected XPD may be more likely to advance along the DNA during melting events. To this end, we examined the position of the helicase on the hairpin prior to each RPA2 melting event and after the duplex reannealed. We identified many (*N* ≈ 400) individual melting events during XPD unwinding and aligned them along the position axis such that each event began at 0 bp before melting occurred (**Fig. 3d**, schematic). We then independently aligned the melting and reannealing transition of each event in time at *t* = *t_melt_* and *t* = *t_anneal_*, respectively (**Fig. 3d**, “before” and “after” schematic). As shown in **Figure 3e,** the resulting probability density of these aligned traces reveals the average position of XPD prior to and following an RPA2 melting event (see **Methods**). After a melting event, we find that the hairpin is most likely to reanneal all the way back to the same position as before the event, indicating that XPD has not moved from its initial position. Importantly, this result shows that there is no significant forward movement of XPD as a result of RPA2 duplex melting, and that XPD does not exploit RPA2’s melting activity to enhance its own unwinding.

**Figure 3f** shows a control in which the alignment process above was repeated at random points during unwinding, each point paired with one *t* = 0.06 s after, corresponding to the time window over which we searched and identified RPA2 melting events (see **Methods**). Again, XPD on average shows no net progress in this amount of time. Comparing **Figure 3e** and **3f** shows that the RPA2-mediated melting events have little to no impact on XPD movement.

### XPD exhibits two distinct types of activity, the fraction of which depends on RPA2

The burst processivities at different [RPA2] (**Fig. 2c**) show that a persistent and significant fraction of bursts exhibit relatively low processivity, ≲25 bp. Bursts with higher processivity extending from 25 to 89 bp also exist at all [RPA2], with their number apparently increasing as [RPA2] increases. An important question is whether there exist any characteristic differences between bursts during which more or less DNA is unwound.

To answer this question, we grouped bursts into two categories: “high-processivity” or “low-processivity” depending on whether or not a threshold of 25 bp was exceeded, this value chosen based on where the tail in the distribution of processivities occurred (**Fig. 2c**). **Figure 4a** shows the unwinding portions of many bursts aligned to begin at the same time (*t* = 0), color coded by [RPA2] and processivity category (light color for low-processivity; dark color for high-processivity; see **Table 1** for number of data traces). When grouped in this manner, some notable features emerge. The fraction of bursts categorized as high processivity increases clearly with RPA2 concentration (**Fig. 4a**) saturating to ~50% at RPA2 concentrations >10 nM (**Fig. 4b**). Furthermore, long stalls (sometimes exceeding tens of seconds in duration) around 10-15 bp are evident at all concentrations and in both high- and low-processivity traces. This stalling behavior has been reported previously (33) and attributed to XPD’s encounter with GC-rich regions—which are more difficult to unwind due to higher base-pairing energy—at this position in the hairpin. All low-processivity bursts exhibit these stalls, after which XPD either backslides to the base of the hairpin or dissociates, ending the burst. However, we have observed XPD continue unwinding past 25 bp after a long stall and be categorized as “high processivity”.

**Figure 4.**
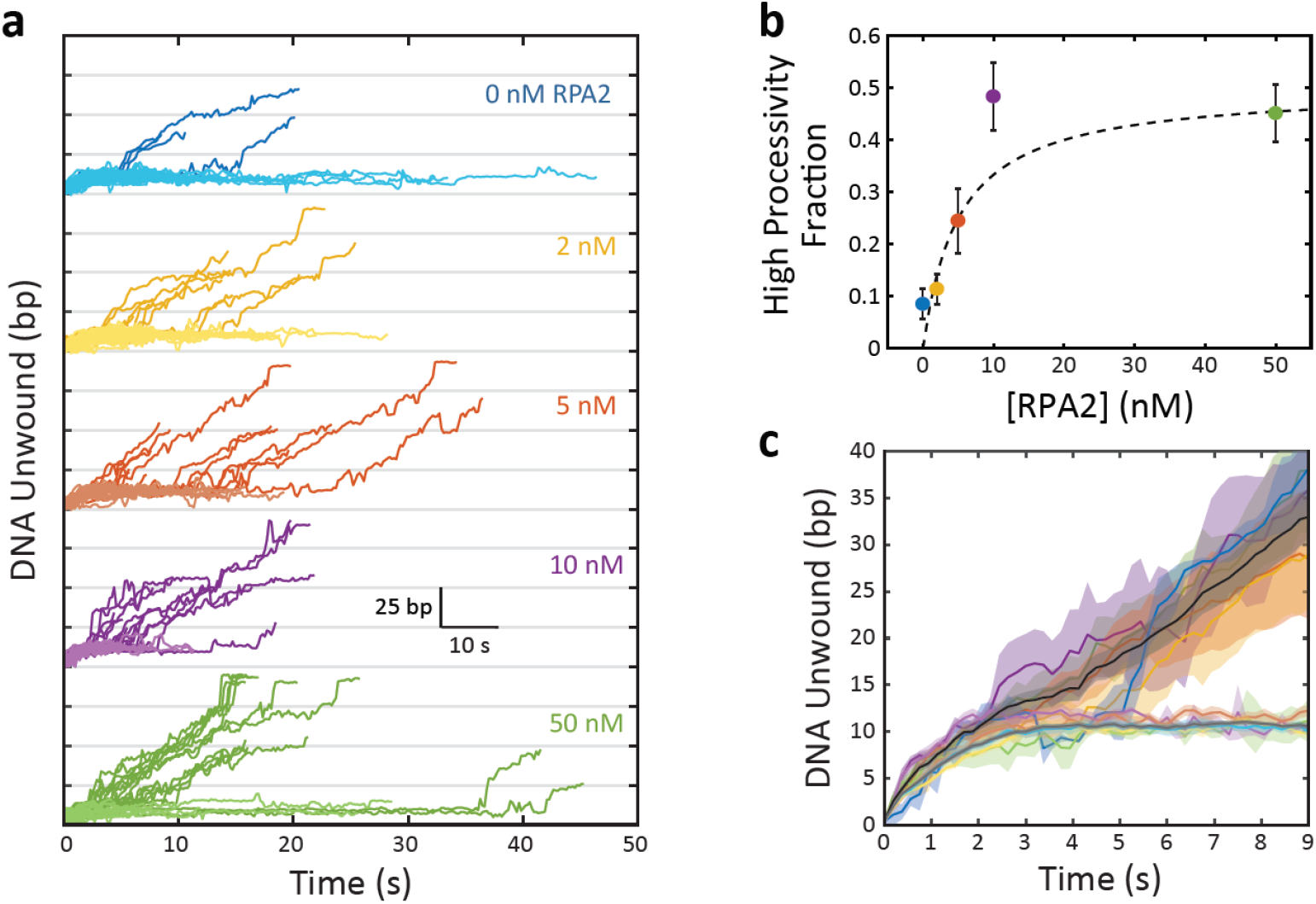
XPD exhibits two burst types, the fraction of which is RPA2 dependent. **a** Plot of XPD unwinding bursts, aligned to start at *t* = 0 and grouped by RPA2 concentration (colored traces). XPD unwinding bursts come in two types: low processivity, never unwinding more than 25 bp, (light colors); and high processivity, unwinding more than 25 bp (dark colors). **b** The fraction of high-processivity bursts (>25 bp) increases with RPA2 concentration. Fit to model described in the text (dashed line; see Methods). **c** Averages of all low-processivity bursts (light colored lines) and all high-processivity bursts (dark colored lines) at each RPA2 concentration. Comparison to averages of low- and high-processivity burst types over all RPA2 concentrations (dark gray and black lines, respectively). Shaded regions represent s.e.m. throughout. Unwinding behavior within each burst category remains the same over all RPA2 concentrations.

**Table 1.** Data statistics.

| Parameter | [RPA2] = 0 | 2 nM | 5 nM | 10 nM | 50 nM |
| --- | --- | --- | --- | --- | --- |
| No. of XPD molec. | 20 | 19 | 27 | 20 | 21 |
| No. of bursts | 94 | 123 | 79 | 60 | 82 |
| No. of bursts/XPD | 4.7 | 6.5 | 2.9 | 3.0 | 3.9 |
| No. of low-proc. bursts | 86 | 109 | 61 | 31 | 45 |
| No. of high-proc. bursts | 8 | 14 | 18 | 29 | 37 |

Importantly, we do not observe significant differences between bursts of the same processivity category obtained at different RPA2 concentrations. **Figure 4c** shows all unwinding bursts for high and low processivity types averaged together at each RPA2 concentration (color coded the same as in **Fig. 4a**) and across all RPA2 concentrations (black and gray lines, respectively). The averaged unwinding traces exhibit the same behavior independent of RPA2 concentration. Low-processivity bursts all display stalls at ~10 bp, whereas high-processivity bursts unwind past this region. Burst-to-burst differences are reflected by the shaded areas and are consistently smaller than the differences between the two processivity categories. This finding is corroborated when analyzing the velocity of XPD. **Figure 4—Figure Supplement 1** shows the average unwinding velocity determined separately for low- and high-processivity bursts at each RPA2 concentration as a function of XPD’s position on the hairpin. While XPD on average slows down near 10 bp due to the high GC content of this section of the hairpin, the velocity of highly-processive bursts is consistently higher than for low-processivity bursts at this position. On the other hand, differences in velocity are insignificant across varying [RPA2] within the same processivity category.

We also find that both low- and high-processivity burst durations remain constant with RPA2 concentration (**Figure 4—Figure Supplement 2**, gray and open data points, respectively). The more processive bursts tend to have longer durations since XPD travels a longer distance on DNA, and processivity increases with [RPA2]. The increase in burst duration observed when combining both types of bursts is explained simply from the increase in the fraction of high-processivity bursts with increasing [RPA2].

Our results show that high-processivity unwinding in the absence of RPA2 is indistinguishable from that in the presence of RPA2 and likewise for low-processivity unwinding. All parameters we quantified by burst in the same processivity category are independent of [RPA2]. Instead [RPA2] increases the probability of high-processivity bursts. These findings suggest that high- and low-processivity unwinding correspond to intrinsic states of XPD, with RPA2 increasing the likelihood of XPD being in its more processive state.

## Discussion

From our results, we propose that XPD can adopt two intrinsic states, one competent for high-processivity unwinding and the other not. Both states are populated independently of RPA2; highly processive unwinding activity can occur in the absence of RPA2 and is indistinguishable from that detected in the presence of RPA2. RPA2 does not assist XPD unwinding by directly destabilizing the duplex, but rather by shifting the equilibrium toward the high-processivity state. In the data we have collected, the two states are effectively hidden, and we can infer their presence only from the processivity exhibited by XPD, i.e. whether XPD unwinds beyond a threshold we have selected as 25 bp.

**Figure 5a** shows a simple kinetic scheme that describes this model quantitatively. XPD can alternate between “low” and “high” processivity states with rate constants *k*_1_ and *k*_-1_. Here, we assume *k*_1_ to be dependent on RPA2 concentration, whereas all other rate constants are independent of RPA2. In the high processivity state, XPD reaches the 25 bp threshold—and is scored as high processivity—at a rate *k*_2_, corresponding to the time XPD takes to unwind 25 bp, which encompasses all forward and backward steps to this position. Finally, we assume that XPD can dissociate from the DNA in the low processivity state with rate constant *k_off_*. Given this kinetic scheme, we derive an expression for the probability, *P*>25(*t*), that a given XPD molecule unwinds >25 bp by time *t* (see **Methods**), which can be compared directly to experimental data (**Fig. 5c**), obtained from the fraction of XPD molecules that have crossed the 25-bp threshold at each time point (**Fig. 5b**).

**Figure 5.**
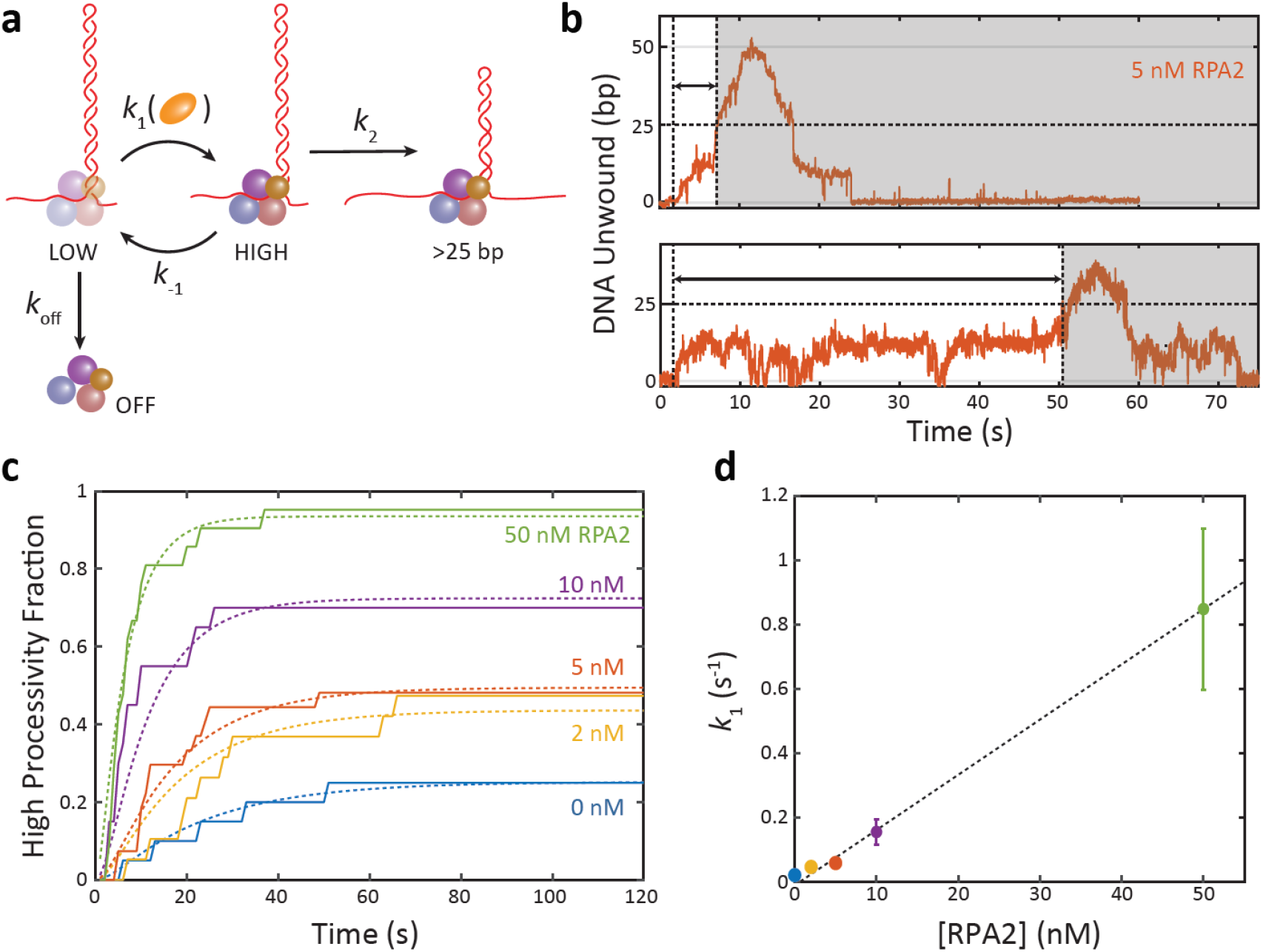
RPA2 activates a high processivity state of XPD. **a** Kinetic model of XPD processivity and the effect of RPA2. XPD can adopt one of two intrinsic states—a LOW and HIGH processivity state. XPD interconverts between both states with rates *k*_1_ and *k*_-1_, and RPA2 shifts the equilibrium toward the HIGH state. XPD can dissociate from DNA only from the LOW state with rate *k_off_*. Only in the HIGH state is XPD able to unwind in excess of 25 bp, which occurs at a rate *k*_2_. Once an XPD molecule unwinds >25 bp, it is scored as being in the HIGH processivity state. **b** Representative traces of XPD exhibiting high processivity unwinding. The time *t*_>25_ denotes the first time at which XPD crosses the 25 bp threshold. **c** Fraction of all XPD molecules that have reached high processivity (>25 bp) after time *t* for each RPA2 concentration. The curves are globally fit to the model in (a) using as parameters the rate constants *k*_1_, *k*_-1_, *k*_2_, and *k_off_*. **d** The rate of entry into the high processivity state, *k_1_*, depends linearly on RPA2 concentration. For more details on the kinetic model see Methods.

**Figure 5c** shows a global fit to our data using different values of *k*_1_ for each RPA2 concentration but a common set of values of *k*_-1_, *k*_2_, and *k_off_*. The parameter values can be found in **Table 2**. (We considered an alternative model in which XPD can dissociate in both its low- and high-processivity states by introducing an additional rate constant 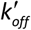 (see **Methods**). The best fit to the data yielded negligible values for 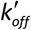, indicating that XPD dissociation occurs in its low processivity state but is unlikely from its high processivity state.) The value obtained for *k_off_* predicts that XPD remains bound to DNA for an average lifetime of 25 s, consistent with observation. A value of *k*_2_ = 0.17 ± 0.02 s^-1^ matches that expected for XPD to unwind 25 bp at an average speed of 4-5 bp/s (33) (**Figure 4—Figure Supplement Fig. 1a**). Finally, we find that *k*_1_ increases linearly with RPA2 concentration (**Fig. 5d**) with a slope of (1.5 ± 0.5) × 10^7^ M^-1^ s^-1^. The linear dependence on [RPA2] suggests that a simple second-order reaction between RPA2 and the XPD-DNA complex shifts the equilibrium to the high processivity state. The intercept of 1-2 × 10^-2^ s^-1^ provides a basal rate of interconversion between the two processivity states in the absence of RPA2. This simple kinetic model can also be compared to our other results. The fraction of high processivity XPD bursts measured vs. RPA2 (**Fig. 4c**) is well fit by the model with no additional fitting parameters (see **Methods**). Likewise, the average burst duration (**Figure 4—Figure Supplement 2**) is well described by the same model (see **Methods**).

**Table 2.** Model fit parameters. Error bars represent 95% confidence intervals.

| Rate const. | Fit value ($s^{-1}$) |
| --- | --- |
| $k_{-1}$ | $0.18 \pm 0.13$ |
| $k_2$ | $0.170 \pm 0.017$ |
| $k_{off}$ | $0.037 \pm 0.019$ |
| $k'_{off}$ | 0 |
| $k_1$, [RPA2] = 0 nM | $0.02 \pm 0.006$ |
| $k_1$, 2 nM | $0.059 \pm 0.017$ |
| $k_1$, 5 nM | $0.065 \pm 0.019$ |
| $k_1$, 10 nM | $0.200 \pm 0.059$ |
| $k_1$, 50 nM | $1.15 \pm 0.39$ |

Although the success of this kinetic model in fitting the data supports our two-state processivity model, it is agnostic to the underlying mechanism of processivity enhancement. We thus must look to other evidence to identify plausible mechanisms. The dependence of the rate *k*_1_ on RPA2 concentration suggests that RPA2 binding to the XPD-DNA complex may activate it for processive unwinding; the linear dependence on [RPA2] rules out mechanisms in which multiple RPA2 bind cooperatively. RPA2 could be binding to DNA, the protein, or both. We have already explored some mechanisms in which RPA2 binding to DNA enhances XPD activity. Our results in **Figure 3** rule out RPA2 melting of DNA as a plausible mechanism. They also disfavor a sequestration mechanism, in which RPA2 binding to ssDNA behind XPD inhibits retrograde motion. By acting as a physical barrier against XPD back-stepping or back-sliding (**Figure 4—Figure Supplement 2a**), two behaviors often exhibited by XPD during unwinding (33), ssDNA sequestration may enhance processivity. Since each burst involves XPD-mediated DNA unwinding followed by rezipping to the base of the hairpin, the duration of a burst provides information on the amount of back-stepping. Bursts with more back-steps are expected to be shorter in duration and vice versa. However, as shown in **Figure 4—Figure Supplement 2b**, we do not observe the mean duration of either the low or high-processivity bursts to increase with RPA2, as predicted by a sequestration mechanism.

Alternately, RPA2 may interact directly with XPD (**Figure 4—Figure Supplement 3**). Direct RPA-helicase interactions have been detected for the Superfamily 2 human helicases WRN, BLM, RECQ, and FANCJ and shown to increase helicase activity (29,30,40–42). While no strong interactions between *Fac*XPD and *Fac*RPA2 have been reported in solution (31), formation of a XPD-RPA2 complex on DNA may be possible. To test such a mechanism, we carried out an experiment in which we allowed a complex of XPD and RPA2 to form on DNA in the absence of ATP, then provided ATP for unwinding. Here, we utilized a different flow chamber configuration in which one stream contained XPD and RPA2 (**Figure 4—Figure Supplement 3b**; red stream) but no ATP, and another stream contained ATP but no protein (blue stream). We placed the DNA hairpin in the protein stream for 40-60 s, then moved it into the ATP stream to observe unwinding. If XPD and RPA2 were to form a complex with higher processivity than XPD alone, we would expect more processive activity at higher RPA2 concentrations as the probability of a complex forming should increase. However, we observed no effect on processivity across RPA2 concentration (**Figure 4—Figure Supplement 3c**); the fraction of bursts that exhibited high processivity (>25 bp) remained constant. The addition of non-hydrolyzable ATP analog ATP-γS to the protein stream had no measureable impact (data not shown). This result indicates that any complex formed would be short-lived, and would not survive the move between the protein and ATP streams, which takes up to ~10 s. The rate constant *k*_-1_ in our kinetic model (**Fig. 5a**) determines the lifetime of the high-processivity state, and its value obtained from a global fit to the data gives a lifetime of ~8 s, consistent with this interpretation. The transience of the interaction may also explain why an XPD molecule may revert to low processivity activity after a high processivity burst and vice versa.

Alternate mechanisms involving transient RPA2 binding to DNA may be more plausible. Recent studies (43) suggest that the displaced (non-translocating) DNA strand may play an inhibitory role for some helicases by binding to secondary sites on the protein surface. Protein binding to the displaced ssDNA could reduce the strand’s inhibitory interaction with the helicase, enhancing unwinding activity. XPD is known to possess a secondary binding site for the (3’) displaced strand located on the HD1 domain, under the ARCH domain (22,44) (**Fig. 1a**). This site is not only important for DNA fork positioning but also plays a regulatory role, likely acting as a throttle to unwinding. In the homologous *T. aquaticus* XPD, a point mutation disrupting this secondary binding site enhances unwinding and force generation (44). Structural analysis of XPD and homologs maps ~5 nt of ssDNA to this secondary site between the DNA fork at the FeS cluster and the entry tunnel into the motor core (33,37,44). Thus, RPA2 binding of 5 nt of ssDNA would disrupt XPD-ssDNA contacts at this secondary site and could similarly activate XPD unwinding. Moreover, previous single-molecule measurements (33) showed that individual XPD exhibit 5-bp backward/forward step pairs (see also **Figure 3—Figure Supplement 2**; blue box), attributed to the transient release/recapture of 5 nt of ssDNA from the secondary site. In the presence of RPA2, we observe the frequency of such events decrease (**Figure 3—Figure Supplement 2**; compare blue box for 0 nM to 50 nm RPA2), consistent with the idea that RPA2 may affect DNA contacts to this regulatory site.

Helicases are likely to encounter other DNA-bound proteins in the cell, and such molecular associations have the potential to regulate helicase activity. Our results provide new insights into the possible mechanisms by which accessory proteins can affect helicases. We show how a single-stranded DNA binding protein can enhance helicase processivity through transient interactions with the helicase-DNA complex which can activate a “processivity switch”. We speculate on the latent conformational states that could be controlled by this switch. The evidence above points to alternate DNA binding configurations (33,44) that regulate unwinding activity, but there may be other possibilities. An intriguing example is the ARCH domain of XPD, which has been shown to interconvert dynamically between two conformations (34). However, while these states are connected to DNA damage recognition (34), their roles in unwinding have yet to be defined. The general mechanism described here for XPD may apply broadly to other helicases. Recent studies of SF2 helicase RecQ (43) suggest a similar inhibitory role of a secondary DNA binding site, which could plausibly be modulated by DNA-binding proteins. Likewise, studies on SF1 helicases UvrD, Rep, and PcrA have shown that alternate conformations of one subdomain strongly affect activity, particularly processivity (39,45). Interactions with protein partners favoring particular subdomain conformations may provide a mechanism for activating processive unwinding (45,46). Identifying the molecular details of the conformational states that produce high and low processivity activity is a rich subject ripe for future investigation.

## Acknowledgements

We thank members of the Chemla and Spies laboratories for scientific discussions. Work in the Chemla lab is supported by National Institutes of Health grant R01 GM120353. Work in the Spies lab is supported by National Institutes of Health grant R35 GM131704.

## Methods

### DNA Construct Synthesis

The hairpin construct was synthesized as described in Qi *et al*. (33). The construct consists of three separate dsDNA fragments ligated together after synthesis and purification (**Figure 1—Figure Supplement 1**): a 1.5-kb ‘Right handle’, an 89-bp ‘Hairpin’ stem capped by a (dT)4 loop, and a 1.5-kb ‘Left Handle’. ‘Right handle’ and ‘Left handle’ are functionalized with a 5’ digoxigenin and biotin, respectively, for binding to anti-digoxigenin and streptavidin coated beads. The hairpin stem sequence used for all experiments was identical to ‘Sequence 1’ used in Ref. (33), which contains a random 49% GC sequence: 5’-GGC TGA TAG CTG AGC GGT CGG TAT TTC AAA AGT CAA CGT ACT GAT CAC GCT GGA TCC TAG AGT CAA CGT ACT GAT CAC GCT GGA TCC TA-3’. The hairpin stem is flanked by a 5’ 10-dT binding site for loading XPD and a 3’ abasic site to prevent XD unwinding into the handle. All oligonucleotides were purchased from Integrated DNA Technologies (Coralville, IA).

### Flow Chamber for Optical Tweezers Measurements

All measurements were carried out in laminar flow chambers described in Whitley *et al*. (36). Chambers were made from NescoFilm (Karlan, Phoenix, AZ) melted between two glass coverslips and laser-engraved with channels for buffers. Chambers for all measurements had three channels (**Figure 1—Figure Supplement 2**). Top and bottom channels contained anti-digoxigenin and DNA-coated streptavidin beads, respectively. Small (100-μm OD) glass capillaries connected the top and bottom channels to the central channel for trapping and allow a controlled flow of beads. Three separate streams converged into the central trapping channel. Because the flow in each stream is laminar, mixing between different buffer streams is minimal and a reasonably sharp boundary between buffer conditions was maintained (**Figure 1—Figure Supplement 2a**, inset). This chamber design allowed moving freely between different streams via motorized sample stage and changing buffer conditions during an experiment, as described below.

### Optical trap measurements

High-resolution dual-trap optical tweezers based on a previously reported design (36,47) were used to study XPD helicase unwinding in the presence of RPA2. The optical traps were calibrated by standard procedures (36,48) and both traps had a typical stiffness of *k* = 0.3 pN/nm in all experiments. All data were acquired using custom LabVIEW software (National Instruments, Austin, TX), and were collected at a 267-Hz sampling rate and boxcar filtered to a lower frequency as indicated in the text. All measurements were carried out at a constant force of 12 pN using a feedback loop to maintain constant force.

*Fac*XPD and *Fac*RPA2 were purified as described previously by Pugh *et al*. (31). Single XPD unwinding experiments were performed in a similar manner to that described in Qi *et al*. (33), with some amendments. The trapping buffer consisted of 100 mM Tris-HCl (pH 7.6), 20 mM NaCl, 2 mM DTT, 3 mM MgCl2, 0.1 mg/mL BSA and oxygen scavenging system (49) (0.5 mg/mL pyranose oxidase [Sigma-Aldrich, St. Louis, MO], 0.1 mg/mL catalase [Sigma-Aldrich, St. Louis, MO], and 0.4% glucose) to increase the lifetime of the DNA tethers (50). To this buffer, varying concentrations of XPD, ATP, ATP-γS, and RPA2 were added. Unless otherwise noted, the following buffers filled the three laminar flow streams in the central trapping channel (**Figure 1—Figure Supplement 2a**): Upper stream, 500 μM ATP + 0-50 nM RPA2; Middle stream, 500 μM ATP-γS + 10 nM Cy3; Lower stream, 60 nM XPD + 500 μM ATP-γS. ATP-γS was used to increase the binding efficiency of XPD to the DNA loading site. The dye molecule Cy3 was added to the middle stream to detect, via fluorescence imaging, the precise locations of the middle stream boundaries. Fluorescence detection of the Cy3 stream was achieved using a confocal microscope incorporated into the optical trap instrument, as described in Refs. (36,47).

During a typical experiment, a single DNA hairpin was tethered between an optically trapped streptavidin-coated bead and an anti-digoxigenin coated bead in the middle stream, in the absence of any protein or ATP (Step 1 in **Figure 1—Figure Supplement 2b** and **2c**). A low force (~2 pN) was applied to the tether, and the traps and tethered DNA were moved by motorized sample stage into the lower stream containing XPD + ATP-γS, where they incubated for ~40-60 s to allow a single XPD to bind to the 10-dT ssDNA loading site but not unwind the hairpin (Step 2). Following incubation, the force on the tether was increased to 12 pN and the tether was moved into the upper stream containing ATP + RPA2 (Step 3). Upon exposure to ATP, an XPD molecule bound at the loading site unwound the hairpin ahead of it, which, at constant force, resulted in an increase in the end-to-end extension of the DNA tether (**Figure 1—Figure Supplement 2c**). Unwinding data were typically collected until the tether broke or XPD dissociated.

### Analysis of DNA hairpins and determining base pairs unwound by XPD

A force-extension curve of each tether was usually taken in the middle stream containing no protein to verify proper synthesis. A properly synthesized hairpin unzips mechanically at an applied force of ~15 pN (**Figure 1—Figure Supplement 1b**), and its force-extension curve is well fit to the following model:

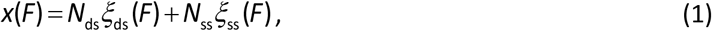

where *ξ*_ds_(*F*) and *ξ*_ss_(*F*) are the extension of 1 bp of dsDNA and 1 nt of ssDNA at a given force, *F*, respectively, and *N*_ds_ and *N*_ss_ are the number of dsDNA base pairs and ssDNA nucleotides, respectively, in the construct. *ξ*_ds_(*F*) and *ξ*_ss_(*F*) are given by the extensible worm-like chain (XWLC) model of elasticity, using the following parameters: persistence length *P_ds_* = 50 nm and *P_ss_* = 1.0 nm, contour length per base pair/nucleotide *h_ds_* = 0.34 nm bp^-1^ and *h_ss_* = 0.59 nm nt^-1^, and stretch modulus *S_ds_* = 1,000 pN and *S_ss_* = 1,000 pN (33,51). For the closed hairpin at forces <15 pN, the model Eq. (1) was used with *N_ds_* = 3,050 bp, corresponding to the sum of the handle lengths, and *N_ss_* = 10 nt, corresponding to the helicase loading site length (**Figure 1—Figure Supplement 1b**, black dashed line). For the open hairpin at forces >15 pN, values *N_ds_* = 3,050 bp and *N_ss_* = 192 nt were used (**Figure 1—Figure Supplement 1b**, gray dashed line).

As a helicase unwinds the tethered DNA hairpin at a constant force, the tether extension increases due to the release of 2 nt of ssDNA for each bp of the hairpin dsDNA unwound. The number of base pairs unwound at each time point was obtained from the relation

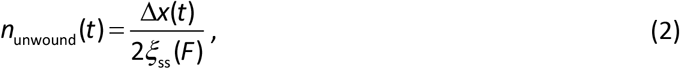

at the force *F* applied, and where Δ*x* is the measured change in extension (in nm).

### Analysis of XPD unwinding bursts

All XPD unwinding time traces were down-sampled from 267 Hz to 89 Hz. Start and end times for each unwinding burst were selected manually using the following criteria: Bursts were defined as periods of significant forward progress by XPD (>5 bp) followed by significant backward motion (>5 bp), excluding behavior obviously attributed to RPA2-mediated transient melting or 5-bp back- and forward step pairs XPD is known to take (33). The processivity (**Fig. 2c**) for each burst was determined from the maximum number of base pairs open between the burst start and end time points, and the burst duration (**Figure 4—Figure Supplement 2**) was determined from the time elapsed between start and end time points.

For the analysis shown in **Fig. 4**, we considered only the unwinding portion of each burst. For highly processive bursts, this segment consisted of the data from the beginning of the burst to the time point at which the maximum number of base pairs was unwound. However, for low-processivity bursts that exhibited extended stalls at ~10 bp, the unwinding endpoint was selected to be the last point above 10 bp. This allowed us to account for extended stalling behavior, while excluding rapid rezipping at the end of the burst. For bursts with processivity less than 10 bp, the unwinding portion of each burst was selected to be the first 75% of the total burst duration to exclude rapid rezipping at the end of the burst.

### Analysis of XPD unwinding speed

To calculate the unwinding speed during each burst, we first smoothed the data to remove any RPA2-mediated transient melting events. Data were smoothed over a 51-point (190-ms) window using the weighted local regression method “rloess” in Matlab’s in-built “smooth” function, which fits the span around each data point to a 2^nd^ degree polynomial and excludes outliers based on a residual analysis. Then, the local velocity was determined over half-overlapping 50-point windows from the slope of a fit of the data to a line. The corresponding times and hairpin positions were also determined from their mean in each window. To create **Figure 4—Figure Supplement 1**, the local velocity for each burst was plotted against hairpin position and averaged over the set of bursts analyzed.

### Analysis of RPA-mediated melting events

RPA2-like melting events on the bare hairpin were identified from short-lived melting of >4 bp of the hairpin (**Figure 3—Figure Supplement 2**). To find RPA2-like events during XPD unwinding, we identified steps in selected XPD unwinding bursts using an algorithm by Kerssemakers *et al*. (52) (**Figure 3—Figure Supplement 2a**). Specifically, we searched for adjacent pairs of steps that met the following criteria: >2 bp forward step (step *n*) followed by >2 bp back-step (step *n*+1) within 5 time points (56 ms) of each other (**Figure 3—Figure Supplement 2b** outlined in red). While false positives are observable even in the absence of RPA2, their probability is low, and the probability increases with RPA2 concentration (**Figure 3—Figure Supplement 2c**), confirming their connection to RPA2 activity.

Each step was then aligned at full bandwidth (267 Hz) to the detected locations of RPA2 melting and reannealing. We used 2D kernel density estimation to visualize the probability density of these aligned traces. In this method, each point in the aligned data was replaced by a kernel function, here a 2D Gaussian. The sum of all kernels provides an excellent approximation to the probability density. The darker color in **Fig. 3e** and **f** indicate higher probability.

### Kinetic model

We defined a model in which XPD can exist in two intrinsic states—a low- and high-processivity state—the latter of which we infer only when XPD unwinds past a ~25 bp threshold. In this model, XPD can alternate between “low” and “high” states (with rates constants *k*_1_ and *k*_-1_) and can also dissociate from DNA from both states (with rates constants *k_off_* and 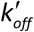, respectively; **Fig. 5a**). The effect of RPA2 increasing the probability of the high-processivity state is captured by the dependence of the rate constant *k*_1_ on [RPA2]. Since the high-processivity state is “hidden”, and unwinding bursts are scored as high processivity only when XPD unwinds >25 bp, the model includes the rate constant *k*_2_ that determines the time XPD takes to pass this threshold.

We used this kinetic model to fit the measured first-passage-time for XPD to unwind past 25 bp. The model in **Figure 5a** is described by a set of master equations for the state probabilities:

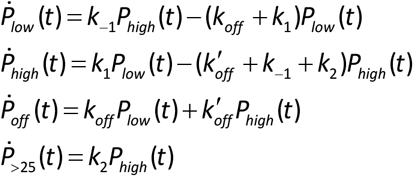

where the subscripts correspond to the four possible states: low processivity (“low”), high processivity (“high”), dissociated (“off”), and past the 25 bp threshold (“>25”). The first passage time is measured from the time XPD is first exposed to RPA2, and XPD is in the low-processivity state. Thus, the initial conditions are given by:

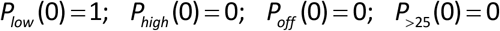

These coupled linear differential equations can be solved to determine all four state probabilities. Here, we provide an expression for *P*_>25_, which is plotted in **Fig. 5c**:

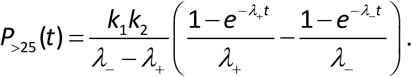

The eigenvalues of the system give the decay constants for the exponentials:

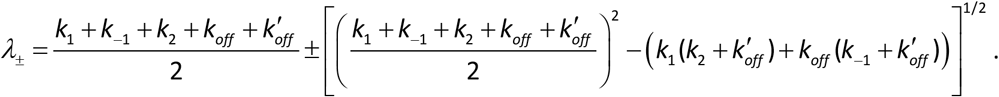

In our model, only *k*_1_ is expected to vary as a function of RPA concentration. By grouping terms that are RPA-concentration dependent vs. independent, *P*_>25_(*t*) can be written in terms of 5 unknown parameters *A, B, C, D*, and *k* related to the 5 rate constants:

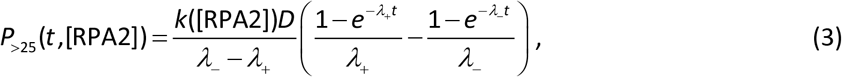

and

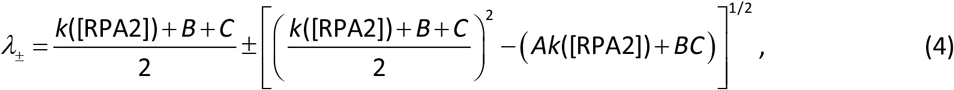

where *k* is written explicitly as a function of [RPA2]. The relationships between the 5 parameters and rate constants are given by:

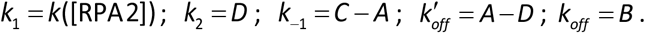

We performed a global fit to our first-passage-time data (**Fig. 5c**) using Eq. (3) and (4) with common values *A, B, C*, and *D* for all RPA2 concentrations but a different value for *k*([RPA2]) at each concentration (**Fig. 5d**). In the data fit displayed in **Figure 5c** (dashed line), we set 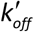 to zero; we found that the best-fit value for 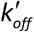 was small with a large error (−0.05 ± 1.23) and provided no significant improvement to the fit. By assuming 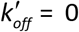, *A* = *D*, reducing our fit to 4 parameters: *A, B, C, k*. Parameter values and uncertainties are listed in **Table 2**.

In addition, we used Eq. (3) to model the RPA2-dependent fraction of high-processivity bursts (**Fig. 4b**) and average burst duration (**Figure 4—Figure Supplement 2**). For the former, data were fit to the expression for *P*_>25_(*t*) evaluated at a time *t* equal to the average low-processivity burst duration *τ_low_* = 7 s. The latter was fit to the following model:

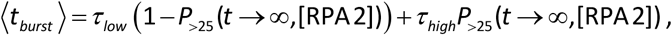

in which the average burst duration is given by the sum of the low-processivity burst duration, *τ_low_*, and the high-processivity burst duration, *τ_high_*, multiplied by the probabilities of being in their respective states at long times (*t* →∞):

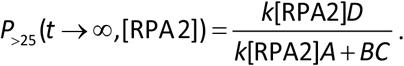

## Figure Supplements

**Figure 1—Figure Supplement 1.**
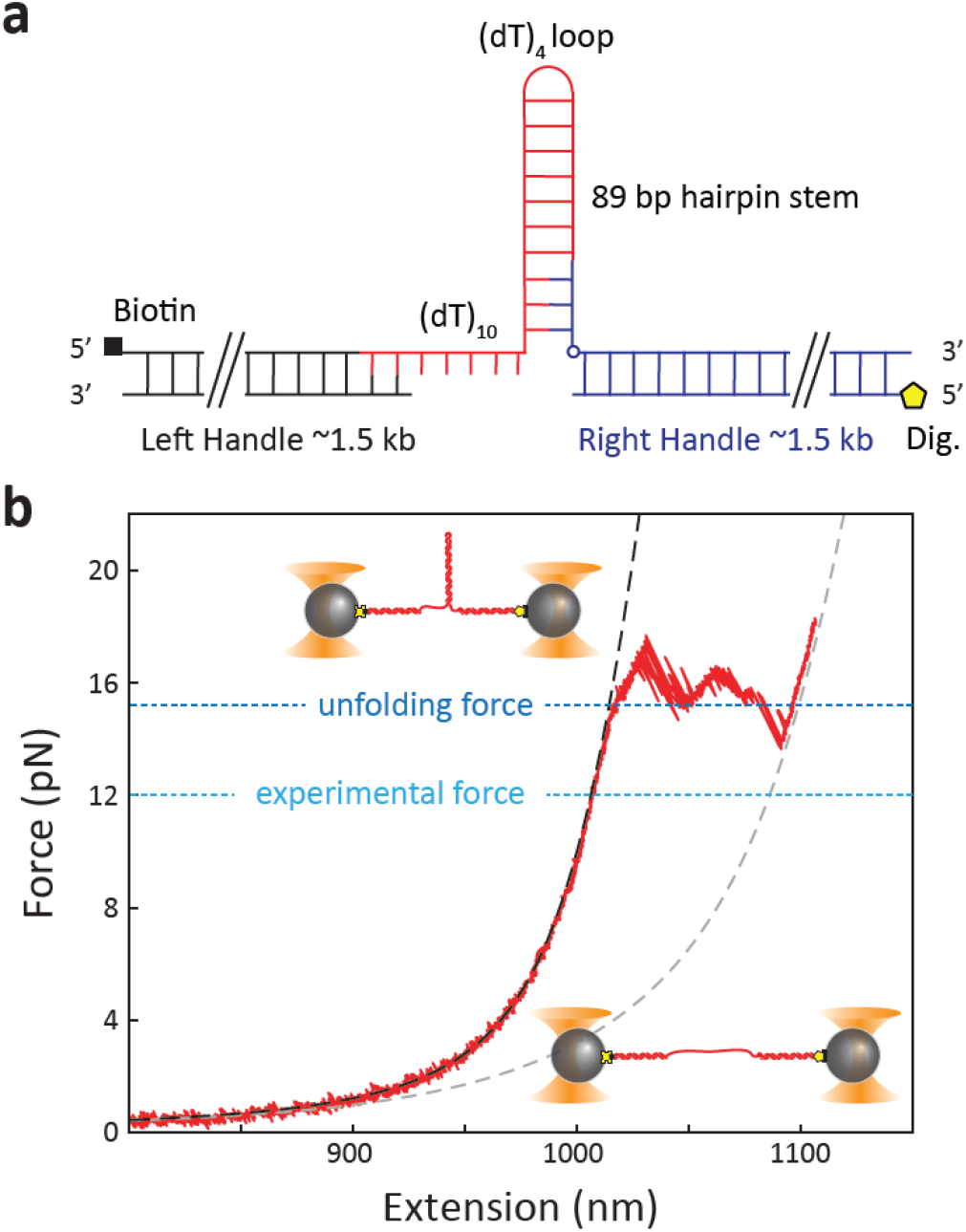
DNA hairpin construct. **a** The hairpin construct consists of three ligated DNA fragments: left (black) and right (blue) dsDNA handles and an 89-bp hairpin that includes a 5’ 10-nt ssDNA protein loading site. The left and right handles are modified with a 5’ biotin (black square) and 5’ digoxigenin (yellow pentagon), respectively, for attachment to beads. **b** Representative force-extension curve of a DNA hairpin. The experimental curve (red solid line) is fit to a model of the closed (black dashed line) and open (gray dashed line) hairpin. (See Methods).

**Figure 1—Figure Supplement 2.**
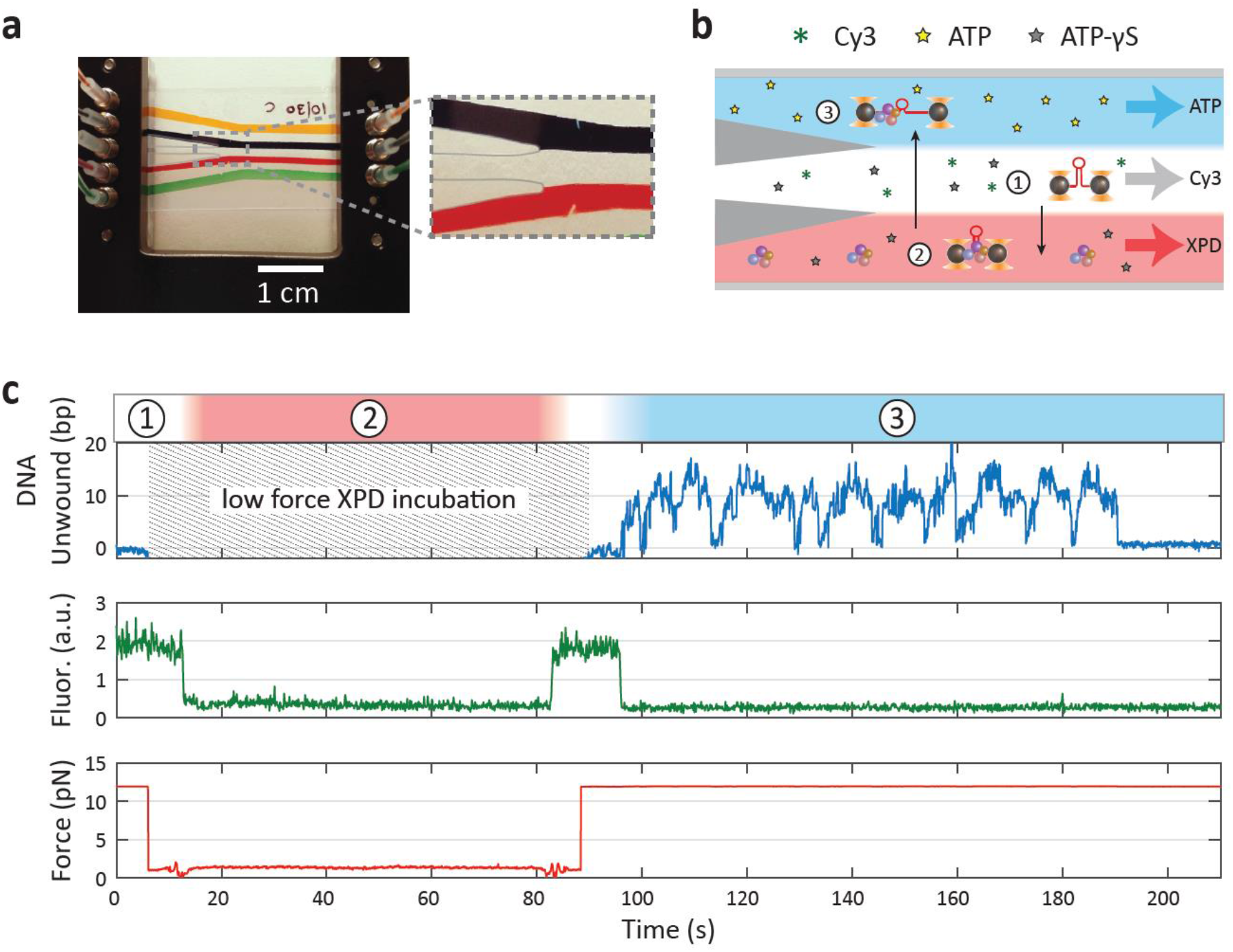
Laminar flow chamber. **a** Photograph of laminar flow chamber. Colored food dye highlights the different flow streams. Parafilm in the central channel is recolored to enhance contrast. The bead channels (yellow and green) are connected via capillaries to the central trapping region (inset). Three streams (dark blue, uncolored, and red) meet in the center channel but mix minimally. **b** Typical experimental protocol. The upper stream of the the central channel is filled with buffer containing 500 μM ATP, the middle stream with 500 μM ATP-γS, and the bottom stream with 60 nM XPD and 500 μM ATP-γS. The dye Cy3 is added to the middle stream for illustrative purposes. A DNA hairpin is tethered between trapped beads in the middle stream (step 1), and its force-extension behavior characterized (see Figure 1—Figure Supplement 1). The trapped beads and DNA are next moved into the XPD stream (step 2) for ~1 min. for a single XPD to bind at low force, ~1-2 pN. Then, they are moved into the ATP stream (step 3) to initiate XPD unwinding, as the force is increased and maintained at 12 pN. **c** Example unwinding trace of a single XPD. Colored regions and numbers above the trace correspond to chamber locations in b. Cy3 fluorescence intensity indicates when the traps are in the middle stream. Unwinding activity begins as soon as XPD enters the ATP stream (*t* = 95 s). XPD unwinds in repetitive “bursts” of activity for ~90 s until it dissociates.

**Figure 3—Figure Supplement 1.**
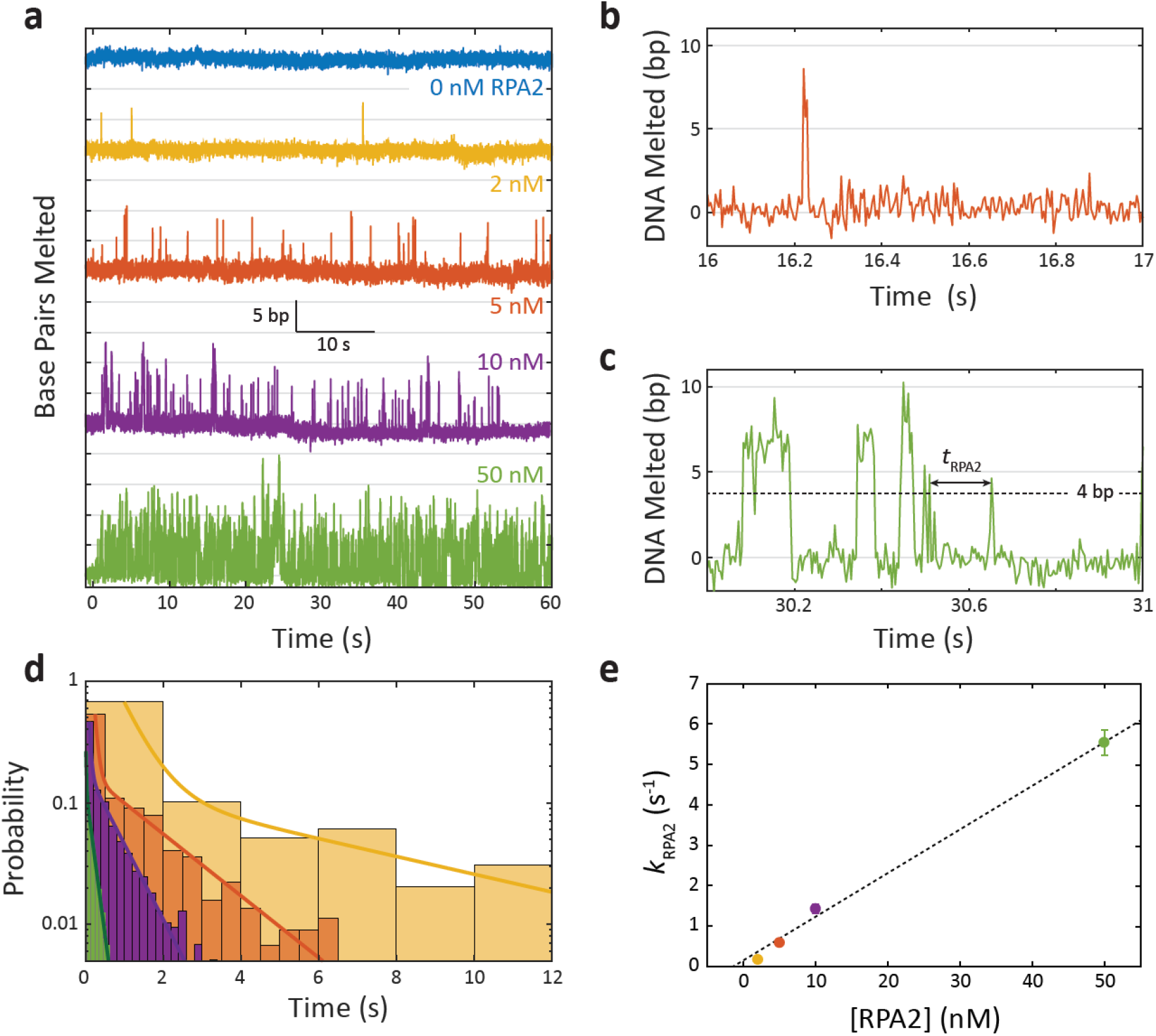
RPA2 transiently melts DNA under force. **a** Representative time traces of RPA2 transiently destabilizing hairpin dsDNA at a constant force (*F* = 12 pN) and at varying RPA2 concentrations. The hairpin is exposed to varying concentrations of RPA2 at *t* = 0. In the absence of RPA2, the hairpin remains stably zipped. In the presence of RPA2, brief (<0.1 s) events consisting of 5-10 bp dsDNA melting followed by rapid reannealing are observed. The frequency of these events increases with RPA2 concentration. **b-c** Selected RPA2 melting events from traces in a with 5 nM and 50 nM RPA2. Melting events are identified at time points where >4 bp of DNA is unwound, and melting kinetics are quantified from the time between events, *t*_RPA2_. **d** Probability distribution of *t*_RPA2_ across RPA2 concentrations (colors same as in a). The distributions are best fit to a bi-exponential with one slow and one fast rate constant. The slow process depends on [RPA2] and corresponds to RPA2 binding and melting the hairpin DNA. The fast process is independent of [RPA2] (data not shown), and attributed to already-bound RPA2 “re-melting” the DNA. **e** The effective RPA2 binding rate, *k*_RPA2_, determined from the slow rate constant in d, increases linearly with RPA2 concentration. Error bars represent s.e.m.

**Figure 3—Figure Supplement 2.**
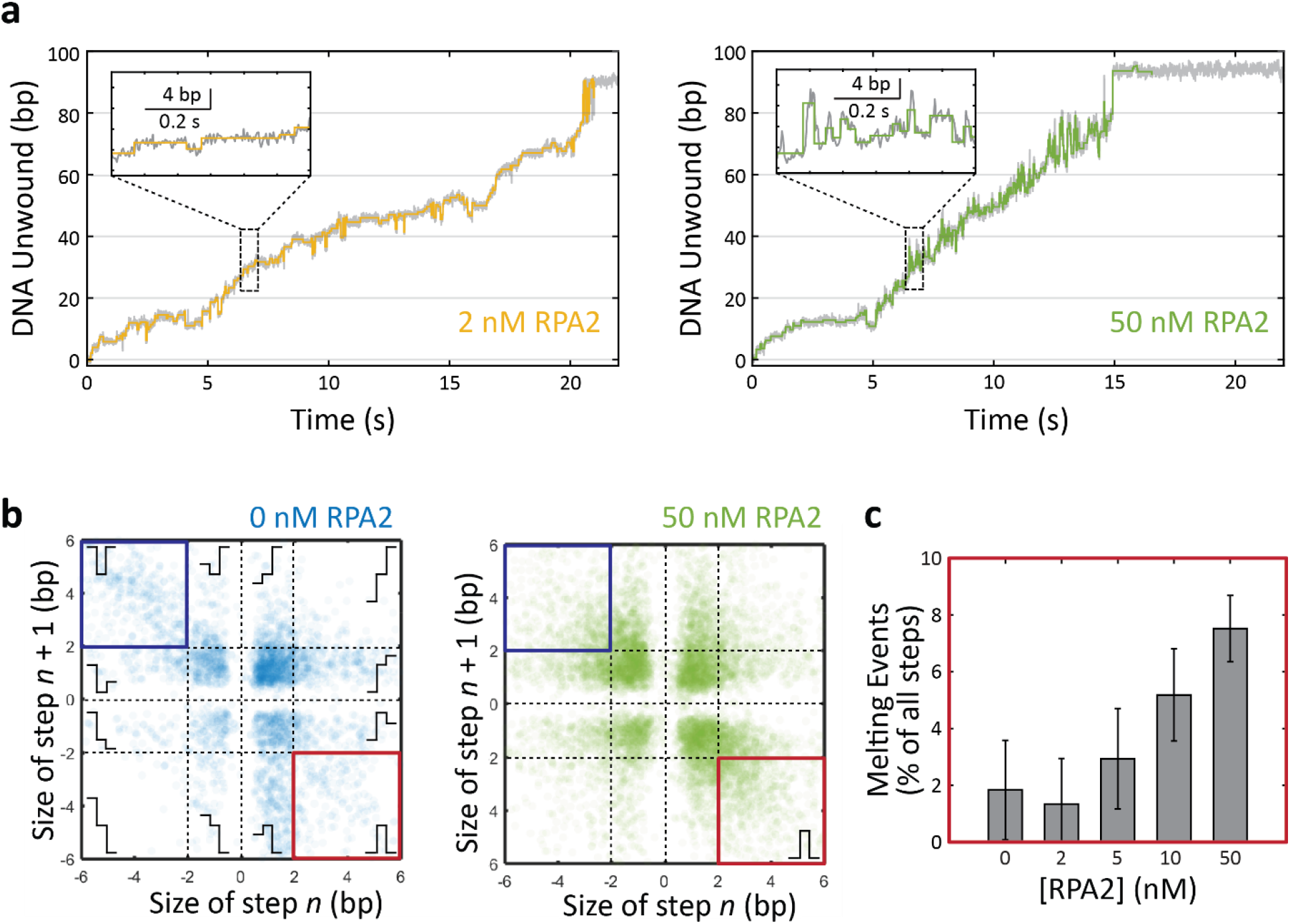
Detecting RPA2 melting events during XPD unwinding. **a** Representative time traces of XPD unwinding in the presence of 2 nM and 50 nM RPA2 (data in gray) and corresponding step-fitting analysis (colored line) used to identify putative RPA2 melting events. Inset: steps fitted (colored lines) to unwinding data (gray). **b** Scatter plot of step size for pairs of consecutive steps for XPD unwinding in 0 and 50 nM RPA2 (blue and green, respectively). Schematics illustrate the step pairs. In the absence of RPA2, small steps (<2 bp) account for the majority of unwinding behavior. In the presence of RPA2, a detectable fraction of step pairs corresponds to a large (>2 bp) forward step followed by a large backward step (red outline; lower right corner). A subset of these events with the appropriate duration (~0.06 s) are selected as RPA2 melting events. Step pairs consisting of a ~5-bp backward step followed by a forward step (dark blue outline; upper left corner) are attributed to DNA dynamics at a secondary, regulatory binding site on XPD. These events are less frequent in the presence of RPA2 (compare 0 nM and 50 nM RPA2; see Discussion). **c** The probability of RPA2 melting events identified in b increases with RPA2 concentration, confirming their connection to RPA2 activity. Some events are detected in the absence of RPA2 due to detection errors. Error bars represent s.e.m.

**Figure 4—Figure Supplement 1.**
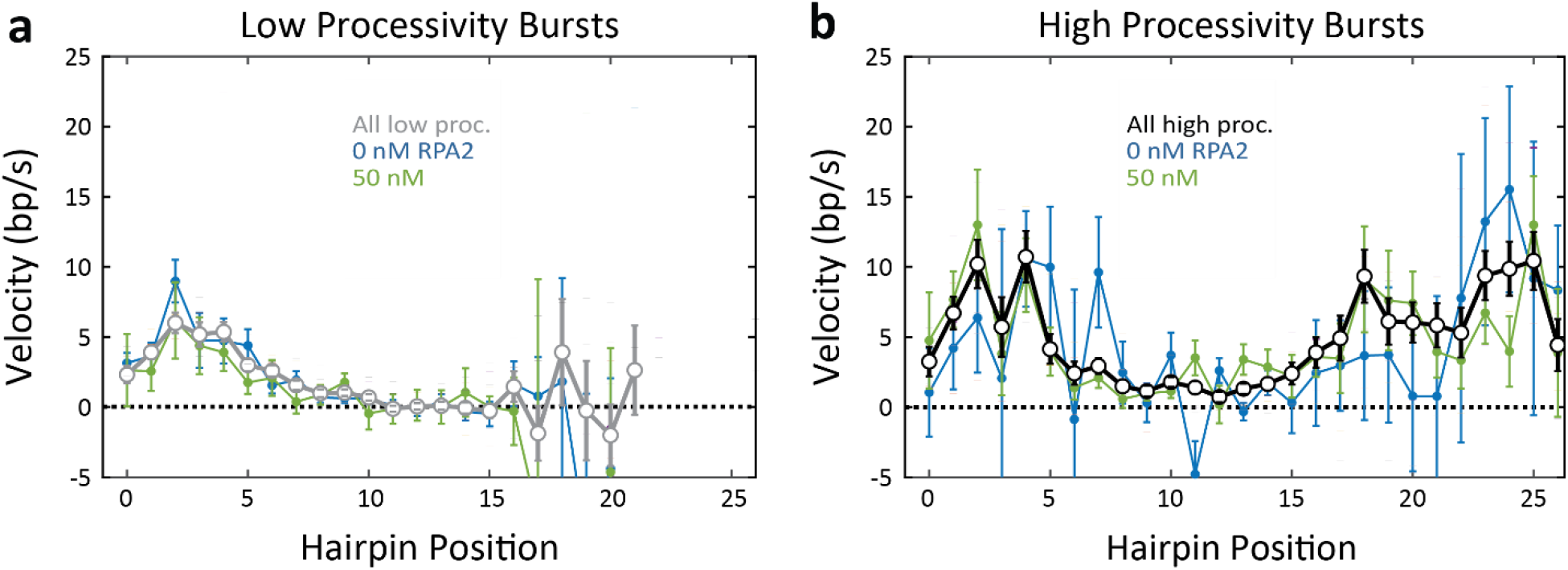
Unwinding velocity varies with processivity type but not RPA2 concentration. **a-b** Average unwinding velocity as a function of hairpin position for low-processivity (a) and high-processivity (b) bursts at two representative RPA2 concentrations, 0 (blue) and 50 nM (green). Velocities averaged over all RPA2 concentration for high- and low-processivity burst types are shown for comparison (black and gray, respectively). Velocity profiles are similar within each processivity type, independent of RPA2 concentration. However, velocity profiles for high processivity bursts differ significantly from those of low processivity bursts, particularly at positions >10 bp. Error bars represent s.e.m.

**Figure 4—Figure Supplement 2.**
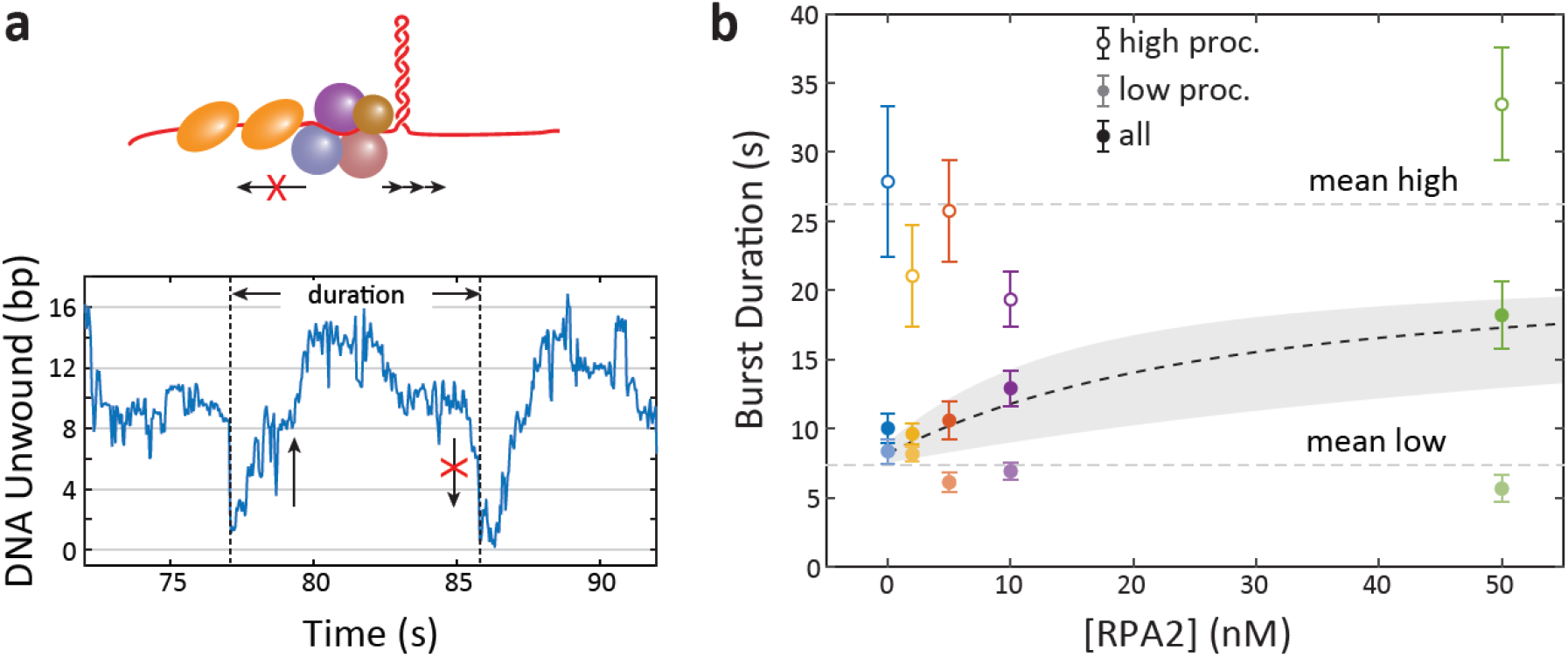
XPD backward motion varies with processivity type but not RPA2 concentration. **a** Schematic representation of the sequestration model, in which RPA2 binding to ssDNA prevents XPD’s backward motion, enhancing unwinding. The model predicts that burst duration should increase with RPA2 concentration, since each burst ends when backward motion takes XPD to the base of the hairpin. **b** Average burst duration for low-processivity (gray circles) and high-processivity (open circles) remains constant with RPA2 concentration. Burst durations averaged over both burst types (black circles) increases with RPA2 due to the increasing fraction of high-processivivity bursts, consistent with the quantitative model (shaded gray region) based on Figure 4b (see Methods).

**Figure 5—Figure Supplement 1.**
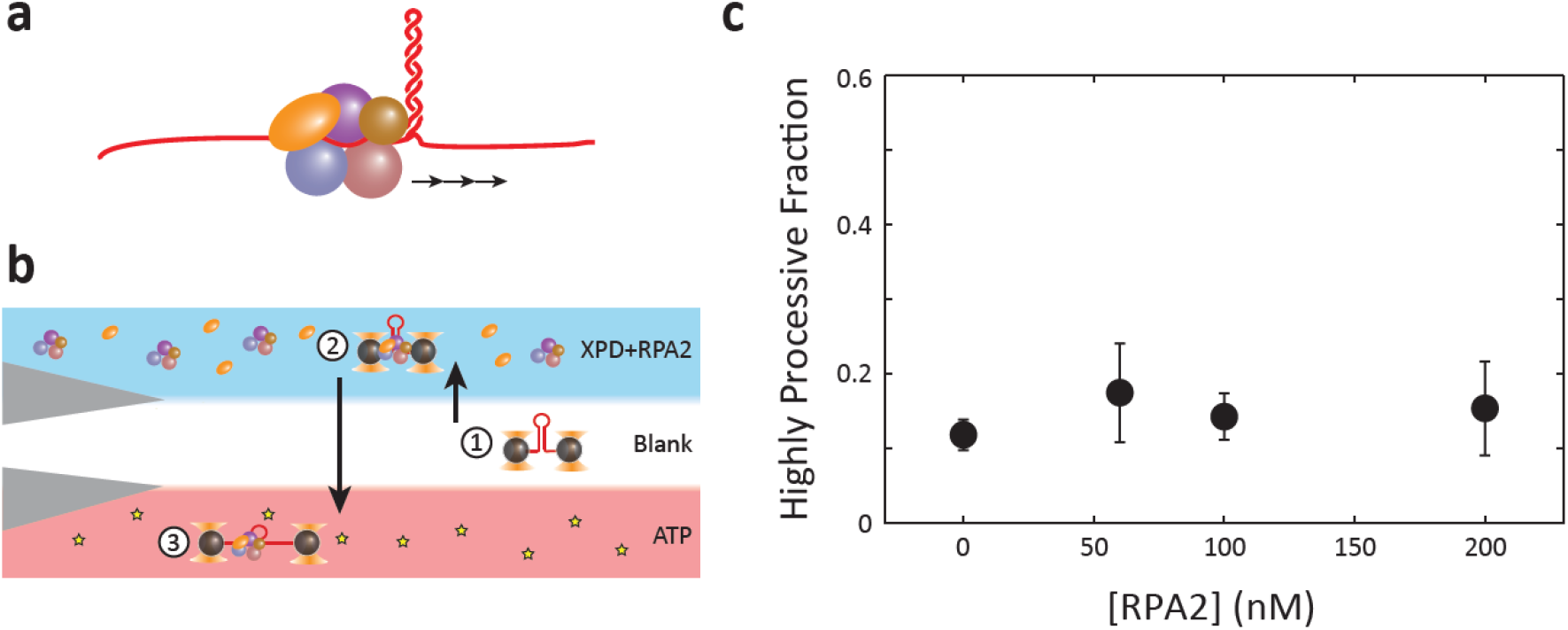
Test of stable complex formation. **a** In the complex formation model, RPA2 forms a complex with XPD that activates it for processive unwinding. **b** Flow chamber configuration for testing the complex formation model. The protein stream contains 60 nM XPD, 500 μM ATP-γS, and 0 – 200 nM RPA2. This configuration allows pre-loading an XPD-RPA2 complex during incubation. Unwinding processivity of the putative complex is measured upon entering into the ATP stream. **c** Although an XPD-RPA2 complex is predicted to be more likely as RPA2 concentration increases, the high processivity fraction remains constant with increasing RPA2 concentration. Error bars represent s.e.m. (*N* = 6 – 43 molecules).

